# IQGAP1 connects phosphoinositide signaling to cytoskeletal reorganization

**DOI:** 10.1101/706465

**Authors:** V. Siddartha Yerramilli, Alonzo H. Ross, Samantha K. Lindberg, Suzanne Scarlata, Arne Gericke

**Affiliations:** From Department of Chemistry and Biochemistry, Worcester Polytechnic Institute, Worcester, MA 01605, USA; Department of Biochemistry and Molecular Pharmacology University of Massachusetts Medical School, Worcester, MA, 01605, USA

**Author notes:** To whom correspondence should be addressed: Department of Chemistry and Biochemistry, Worcester Polytechnic Institute, Worcester, MA 01605, USA.

## Abstract

IQGAP1 is a multi-domain protein that acts as a scaffold for multiple signaling pathways. IQGAP1 generates the lipid messenger PI(3,4,5)P_3_ by scaffolding the phosphoinositide kinases PIPKIs and PI3K. The dynamics of this scaffolding protein complex in intact, living cells are unknown. Here, we delineate the role of IQGAP1 in PI(3,4,5)P_3_-mediated signaling in live cells under basal and stimulated conditions using fluorescence lifetime imaging microscopy. We demonstrate that IQGAP1 interacts strongly with PIPKIγ at intracellular entities and on the plasma membrane, and scaffolds PI3K and PIPKIγ in response to physiological changes. Additionally, we show that IQGAP1 scaffolds phosphoinositides with PI3K, PIPKIγ and EGFR, and forms clusters upon cell stimulation with epidermal growth factor. Importantly, we show that IQGAP1 connects PI(3,4,5)P_3_-mediated signaling and cytoskeletal signaling pathways by binding PIPKIγ in proximity of the cytoskeletal proteins talin and Cdc42. Our results support a model in which IQGAP1 mediates crosstalk between phosphoinositide signaling and the cytoskeleton to promote directed cell movement.

## Introduction

Cell migration is a dynamic process that involves assembly and disassembly of the cell cytoskeleton and focal adhesions. It requires co-ordination between phosphoinositide localization and turnover and cytoskeletal proteins. Enhanced cell migration and dysregulation of phosphoinositide lipids are considered hallmarks of several diseases including cancer [1, 2]. Cell migration requires co-ordination between phosphoinositide lipid signaling pathways and associations with cytoskeletal proteins. In addition to cell migration, phosphoinositides (PIPs) regulate many essential cellular processes by acting as secondary messengers, usually in the form of PI(4,5)P_2_ (phosphatidylinositol-4,5-bisphosphate) and/or PI(3,4,5)P_3_ (PI 3,4,5-triphosphate). Although the overall cellular concentration of PI(4,5)P_2_ is relatively constant, it accumulates at leading edges; these protrusive structures are characterized by intense actin polymerization that is essential for cell polarity [3, 4]. Changes in local PI(4,5)P_2_ concentrations are controlled by redistribution and activation of PI(4,5)P_2_ generating enzymes, such as PI(4)P specific-phosphatidylinositol phosphate kinases (PIPKIs) [5]. PI(4,5)P_2_ can be phosphorylated by class I phosphoinositide 3-kinases (PI3Ks) to generate PI(3,4,5)P_3_ that also localizes to the leading edges of migrating cells [6, 7].

PI(4,5)P_2_ and PI(3,4,5)P_3_ bind proteins that regulate cytoskeletal reorganization at the membrane. These cytoskeletal regulatory proteins include small GTPases such as Cdc42, and several actin regulatory proteins [8–10]. Cdc42 is a major initiator of the formation of the leading-edge, which is a site of actin polymerization and cell polarity. PI(4,5)P_2_ also induces conformational changes in focal adhesion proteins that anchor the actin cytoskeleton to the plasma membrane. By differentially binding to these actin regulatory proteins and focal adhesion proteins, PI(4,5)P_2_ and PI(3,4,5)P_3_ ensure that actin polymerization occurs at the leading edge of migrating cells [3, 11–13].

IQGAP1 (IQ-domain containing Ras GTPase Activating Protein 1) is a large multifunctional protein that facilitates a range of distinct signaling pathways [14], by interacting directly or indirectly with dozens of proteins, thereby affecting key functions of living cells, including proliferation, cell adhesion, motility and metabolism [14, 5]. Recent studies indicate that IQGAP1 associates strongly with multiple PIPKIs, including PIPKIα and PIPKIγ isoforms [16]. In particular, IQGAP1 and PIPK isoforms localize to specific sites on the plasma membrane that include the leading edge of migrating cells, cell-cell contact sites, intracellular membranes and focal adhesions where they share many binding partners, including actin and its regulators. PIPK isoforms are also thought to regulate PI(4,5)P_2_’s associations with actin regulatory proteins and focal adhesion proteins [5]. PI(4,5)P_2_ and PI(3,4,5)P_3_ bind to IQGAP1 at distinct sites as demonstrated by in-vitro studies [16, 7].

IQGAP1’s association with PIPKIα and PI3K but not PIPKIγ are enhanced by various agonists like epidermal growth factor (EGF). IQGAP1 acts as a scaffold for EGF-mediated activation of PI3K/Akt1 and Erk1 signaling pathways [17, 8]. The activation of PI3K/Akt signaling pathway in response to EGF is facilitated by IQGAP1-PIPKI interactions [19] and may play a role in the generation of PI(3,4,5)P_3_ [17]. These results suggest that IQGAP1 promotes the generation of PI(3,4,5)P_3_ by scaffolding the kinases that maintain phosphoinositide pools on the plasma membrane.

Since IQGAP1 lacks catalytic activity [20], it likely plays its signaling role as a scaffold [14]. Even though many IQGAP1 binding partners have been identified, several key questions remain to be answered: First, considering the promiscuity of IQGAP1 with respect to binding partners and signaling pathways, what factors favor scaffolding of one pathway over another? Second, what are the cellular sites where IQGAP1 scaffolds the respective pathways? Third, what is the sequence of scaffolding of various pathways’ components (concurrent vs. successive)? Fourth, is IQGAP1 a scaffold that facilitates crosstalk between different signaling pathways?

IQGAP1 has been studied by biochemical methods using recombinant proteins, but its dynamic interactions in living cells and its ability to coordinate these different signaling pathways in time and space requires further study. Here we use advanced fluorescence methods to show that IQGAP1 integrates PI(3,4,5)P_2_ mediated signaling with cell migration through direct physical interactions between PIPKIγ and PI3K, and the cytoskeletal proteins, talin and Cdc42. Our work shows that IQGAP1 provides a scaffold that allows different signaling pathways to converge to regulate cytoskeletal rearrangements in response to extracellular stimuli.

## Results

### IQGAP1 associates with PIPKIγ

Interactions between IQGAP1 and PIPKIγ occur primarily at the plasma membrane [16], where PIPKIγ may enhance IQGAP1 binding to the plasma membrane (4). This result was corroborated by confocal imaging of HeLa cells expressing both eGFP-IQGAP1 and dsRed-PIPKIγ, which showed a high degree of colocalization **(Fig. 1A)**. It is thought that this PIPKI binding to IQGAP1 causes a conformational change that allows for interactions with additional partners such as PI(4,5)P_2_ [15]. Here we characterize the interactions between IQGAP1 and different phosphoinositide kinases using FLIM-FRET (see *Experimental Procedures*). This method measures the decrease of FRET donor’s lifetime due to transfer to a FRET acceptor. Given the large size of the tagged proteins, any observed changes in donor lifetime in the presence of the proteins tagged with an acceptor can be attributed to changes in physical interactions caused by the direct protein-protein interactions. The FLIM images show the lifetime of eGFP-IQGAP1 in each pixel of the image and these raw values are directly plotted on the phasor plot, i.e., the phasor plot is a true representation of the raw lifetime data without any modifications. In HeLa cells expressing only eGFP-IQGAP1, eGFP-IQGAP1 is found in the cytosol and along the plasma membrane **(Fig. 1B, left)**. The eGFP lifetimes exhibit a largely homogenous population as indicated by a single circular spot on the arc of the phasor plot **(Fig. 1B, middle)**, analogous to cells expressing only free eGFP **(Fig. S1A-B** in appendix). The pixels underlying this spot can be seen as a uniform lifetime across the entire cell except for the nucleus whrer IQGAP1 does not localize **(Fig. 1B, right)**. In HeLa cells, we find that when we co-express PIPKIγ tagged with a FRET acceptor, the localization of eGFP-IQGAP1 shifts towards the plasma membrane (**Fig 1A** and **Fig. 1C)**. The average eGFP-IQGAP1 lifetime decreases by ~10% due to FRET to dsRed-PIPKIγ compared to cells that express only eGFP **(Fig. S2A-D** in the appendix). This decrease in fluorescence lifetime is significantly greater than the ~1-3% decrease in lifetime caused by non-specific binding and indicates an interaction between IQGAP1 and PIPKIγ **(Fig. S1B** in the appendix).

**Figure 1:**
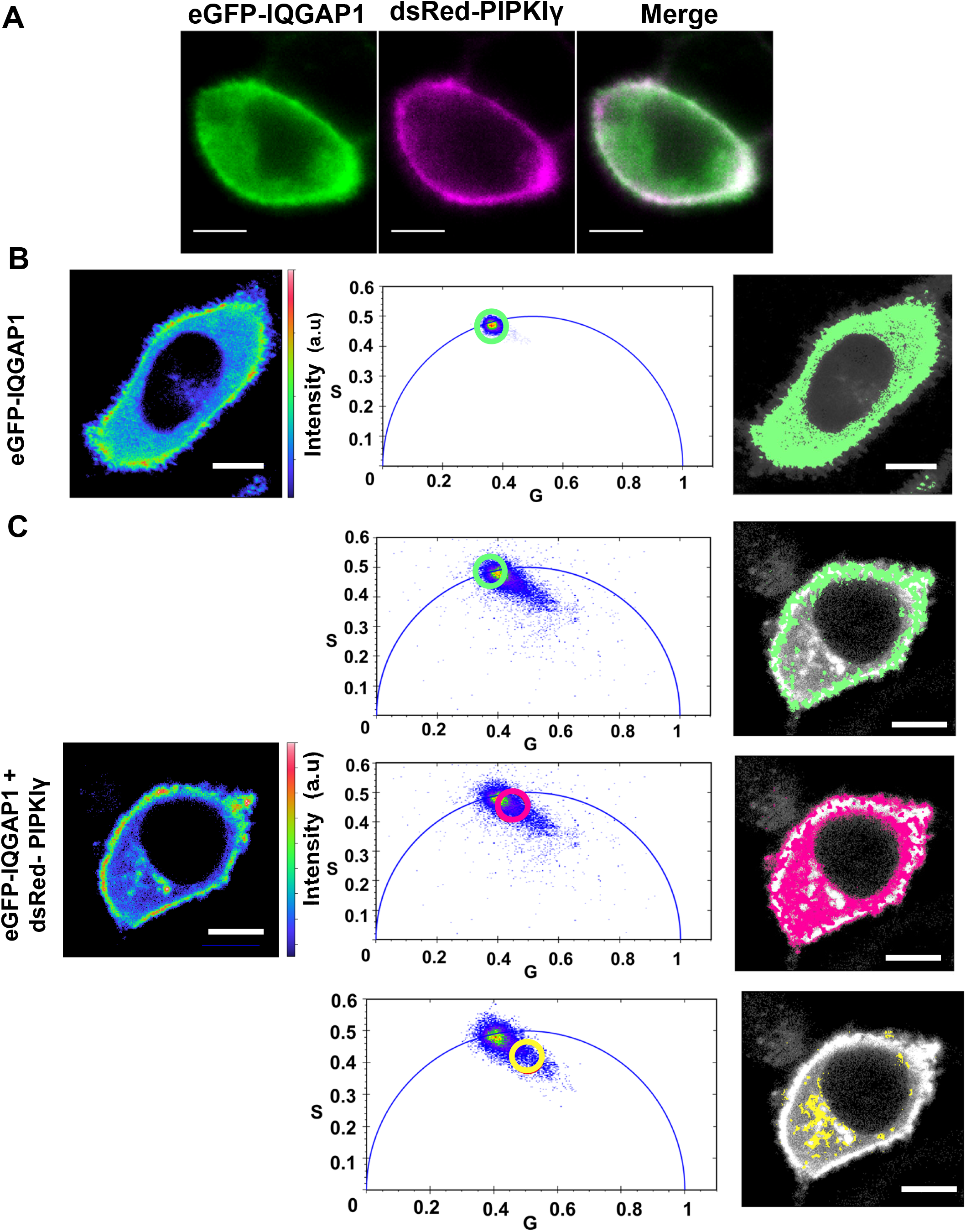
IQGAP1 strongly associates with PIPKIγ: A) A representative confocal microscopy image showing colocalization between eGFP-IQGAP1 and dsRed-PIPKIγ in a HeLa cells B) A false color image of a representative cell (left) shows that eGFP-IQGAP1 expressed in HeLa cells has a broad distribution across the cell except the nucleus. Plotting the raw lifetime values of each pixel in the image as a phasor plot (described in *Experimental Methods*, middle) shows that eGFP-IQGAP1 exhibits a single fluorescent lifetime indicated by a homogenous population on the phasor arc. The pixels included in the green circle of the phasor plot (lifetime center= 2.56 ns) are false colored green and overlaid on a grayscale image of the cell (right), leading to the conclusion that the eGFP-IQGAP1 fluorescence lifetime is similar throughout the cell. The green pixels completely cover the gray of the cell. C) When co-expressed with dsRed-PIPKIγ, the representative cell image (left) shows that IQGAP1 is primarily localized to the membrane. The phasor plots (center panels), show distinct eGFP-IQGAP1 lifetime populations both on the arc and inside the arc due to reduced lifetimes caused by FRET. The distinct regions on the phasor plots are highlighted by a green circle indicating higher (non-FRET) lifetimes (lifetime center = 2.55 ns, top) or by magenta and yellow circle indicating shortened lifetimes (magenta lifetime center= 2.00 ns, yellow lifetime center= 1.60 ns). The pixels underlying these circles are false colored and overlaid on grayscale cell images (right panels).

In phasor plots, FRET us seen by a shift in lifetimes inside the arc giving the data a comet-like appearances. The points inside the phasor arc corresponds to a lifetime with reduced value caused by FRET. The eGFP-IQGAP1 lifetimes in these cells have a ‘comet’-like distribution on the phasor plot that allows us to distinguish distinct FRET and non-FRET pixel subpopulations **(Fig. 1C,** center panels). When these lifetimes (identified in the phasor plot by three circles) are visualized in the whole cell image **(Fig. 1C,** right panels), we see the lower lifetime FRET populations at the plasma membrane as well as other regions in the cytosol indicating the subcellular locations of IQGAP1-PIPKIγ interactions **(Fig. 1C,** right bottom),. It is possible that these cytosolic interactions occur in endosomes and/or lysosomes as IQGAP1 is known to localize to endosomes [21]. A similar interaction between IQGAP1 and PIPKIγ is also seen in NIH 3T3 fibroblasts and a liver cancer cell line HepG2 **(Fig. S2C)**. The interaction between IQGAP1 and PIPKIγ is not perturbed by either the PI3K activator EGF that results in generation of PI(3,4,5)P_3_ or by the PI3K inhibitor LY294002 **(Fig. S2B)**.

### IQGAP1 mediates PIPKIγ association with PI3K

IQGAP1 has been shown to co-localize with PI3K subunits (p85, p110β and p110α) [17], which we corroborated using mass spectrometry **(Table 1)** and immunofluorescence **(Fig. S3B-C,** appendix). A previous study also showed that PIPKIα co-immunoprecipitated with the PI3K p110α subunit in the presence of IQGAP1, even though PIPKIγ and PI3K are not thought to bind directly to each other (12). Here, we observe FRET between dsRed-PIPKIγ with both emGFP-PI3K p110α and eYFP-PI3K p85, indicating an association between PIPKIγ and PI3K in HeLa cells. The interaction between the PI3K p85 subunit and PIPKIγ increases upon stimulation with EGF as demonstrated by a statistically significant decrease in lifetime **(Fig. 2A, B)**, while the FRET between PI3K p110α and PIPKIγ is not affected by EGF. A heterogeneous population is seen on the phasor plot (**Fig. 2A**, center panels). Lifetime imaging shows that FRET occurs in the plasma membrane region as well as some intracellular structures (**Fig. 2A**, right bottom). Because the two kinases do not bind to each other, it is likely that this interaction is due to scaffolding mediated by IQGAP1. We tested this idea by down-regulating IQGAP1 in HeLa cells, which resulted in about 66+/-20% decrease in IQGAP1 protein levels. Upon treatment with IQGAP1 siRNA, the interactions between PIPKIγ and PI3K subunits p85 and p110α are significantly attenuated upon EGF stimulation **(Fig. 2B)**. Meanwhile, treatment with the PI3K inhibitor LY294002 reduced this interaction between dsRed-PIPKIγ and PI3K p110α subunit but not the p85 subunit **(Fig. S3D)**. These results indicate that IQGAP1’s scaffolding of PIPKIγ and PI3K is essential for robust response to EGF that involve the generation of PI(3,4,5)P_3_.

**Figure 2:**
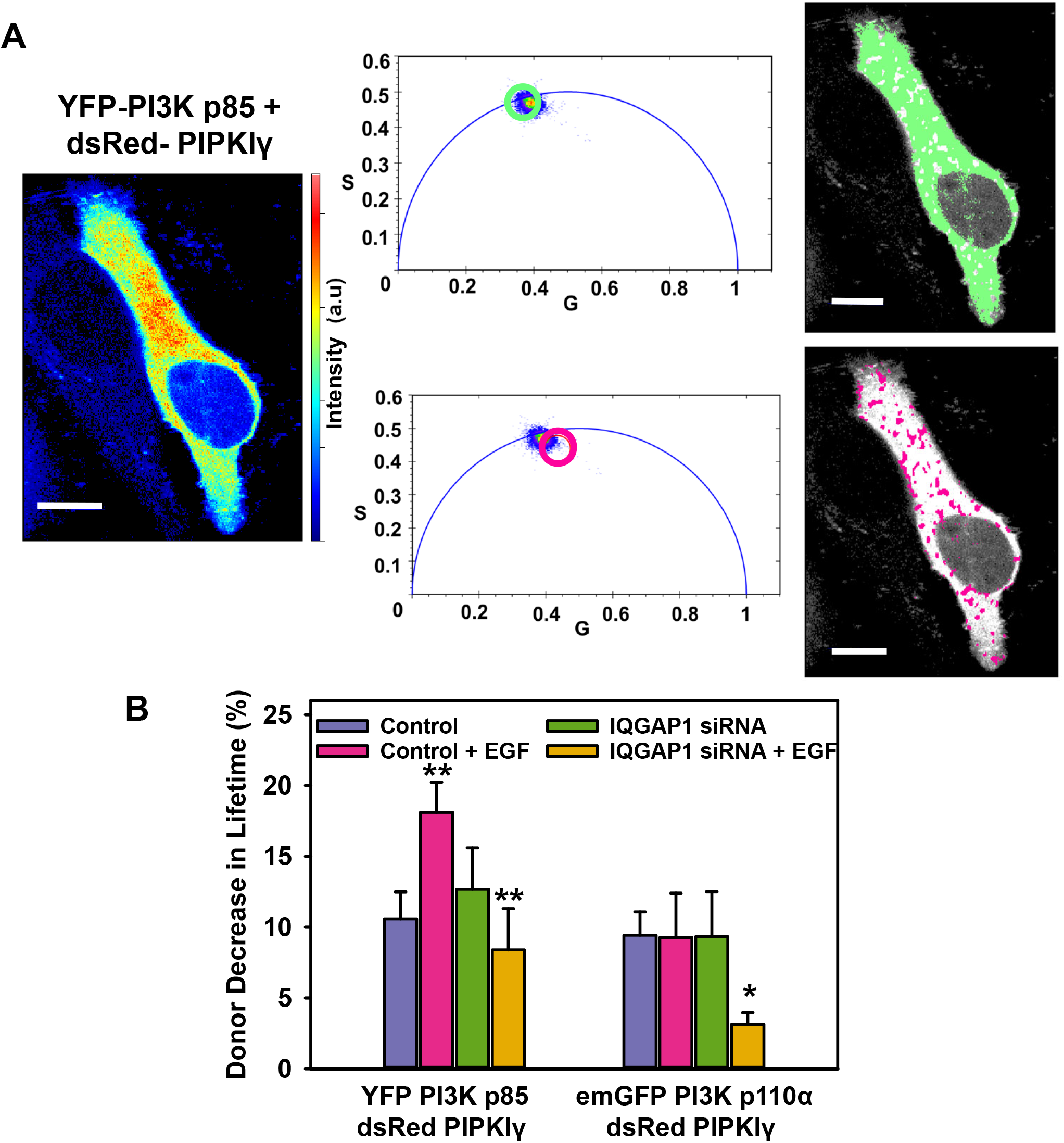
IQGAP1 mediates PIPKIγ / PI3K association: A) A representative cell image shows the eYFP fluorescence of HeLa cells expressing both dsRed-PIPKIγ and eYFP-PI3K p85 (left). eYFP lifetimes for some pixels are inside the phasor arc due to reduced lifetime caused by FRET (center). The distinct regions on the phasor plots are highlighted by a green circle indicating higher (non-FRET) lifetimes (lifetime center = 2.55 ns) and a magenta circle indicating shortened lifetimes (lifetime center= 2.00 ns). The pixels underlying these circles are false colored and overlaid on grayscale cell images (right panels). D) EGF treatment decreases the fluorescence lifetime of HeLa cells expressing eYFP-PI3K p85-dsRed-PIPKIγ-but not emGFP PI3K p110α-dsRed-PIPKIγ, indicating the EGF enhances the interactions of PIPKIγ with p85 but not p110α. Furthermore, suppressing IQGAP1 levels with siRNA decreased the PIPKIγ-p85 and PIPKIγ-p110α interactions indicating that IQGAP1 mediates these interactions. Scrambled (non-specific) siRNA treatment is used for the control groups. (n≥5 measured in at least two independent experiments, * = p ≤0.011, **=p<0.001 from scrambled siRNA alone). In both cases, lifetimes of the control and IQGAP1 siRNA groups are significantly different from eYFP-p85 alone and from emGFP-p110α alone (p<0.001). Error bars denote standard deviation. Scale bar= 10μm

**Table 1-.** Protein partners of IQGAP1 involved in cytoskeletal reogranization: Proteins bound to endogenous IQGAP1 in HeLa cells that were characterized to be involved in actin remodeling and/or being a part of the focal adhesions detected from a coimmunoprecipitation study using anti-IQGAP1antibody using mass spectrometry. The mass spectrometry data was analyzed using the Database for Annotation, Visualization and Integrated Discovery (DAVID)

| Protein name | Previously reported interaction with: |  |  |
| --- | --- | --- | --- |
|  | PI(4,5)P2 | PIPKI | PI(3,4,5)P3 |
| **Kinases and Phosphatases linked to cytoskeleton** |  |  |  |
| PI3K (IQGAP1 pulled down with anti-PI3K 110α) | Y | N | Y |
| Akt1 |  |  | Y |
| Protein phosphatase 1 catalytic subunit alpha(PPP1CA)(PPP1CB) | Y |  |  |
| Mitogen-activated protein kinase 1(MAPK1) |  | Y |  |
| Mitogen-activated protein kinase kinase 1(MAP2K1) |  |  |  |
| Mitogen-activated protein kinase kinase 2(MAP2K2) |  |  |  |
| Integrin linked kinase(ILK) |  |  |  |
| Rho associated coiled-coil containing protein kinase 2(ROCK2) |  |  |  |
| **Small GTPase related proteins** |  |  |  |
| Rho/Rac guanine nucleotide exchange factor 2(ARHGEF2) | Y |  |  |
| **cell division cycle 42(CDC42)** |  | N |  |
| ras homolog family member A(RHOA) |  | Y |  |
| ras-related C3 botulinum toxin substrate 1 (rho family, small GTP binding protein Rac1)(RAC1) | Y | Y |  |
| **Cytoskeleton** |  |  |  |
| **actin beta(ACTB)** | Y | Y |  |
| actin gamma 1(ACTG1) |  |  |  |
| actin related protein 2/3 complex subunit 1B(ARPC1B) |  |  |  |
| actin related protein 2/3 complex subunit 2-5(ARPC2-5) | Y |  |  |
| actinin alpha 1(ACTN1) | Y |  |  |
| actinin alpha 4(ACTN4) | Y |  |  |
| annexin A1,5,6(ANXA1,5,6) | Y |  |  |
| cortactin(CTTN) | Y |  |  |
| myosin heavy chain 9(MYH9) |  |  |  |
| plectin(PLEC) |  |  |  |
| tropomyosin 4(TPM4) |  |  |  |
| vasodilator-stimulated phosphoprotein(VASP) | Y |  |  |
| tensin 3(TNS3) |  |  |  |
| **Focal Adhesion** |  |  |  |
| AHNAK nucleoprotein(AHNAK) |  |  |  |
| cofilin 1(CFL1) | Y |  |  |
| ezrin(EZR) | Y |  |  |
| filamin A, B(FLNA/FLNB) | Y |  |  |
| gelsolin(GSN) | Y |  |  |
| junction plakoglobin(JUP) |  |  |  |
| moesin(MSN) | Y |  |  |
| parvin alpha, beta(PARVA)(PARVB) |  |  |  |
| profilin 1(PFN1) | Y |  |  |
| radixin(RDX) | Y |  |  |
| syndecan binding protein(SDCBP) |  |  |  |
| **talin 1(TLN1)** | **Y** | **Y** |  |
| testin LIM domain protein(TES) |  |  |  |
| vimentin(VIM) |  |  |  |
| vinculin(VCL) | Y |  |  |
| zyxin(ZYX) | Y |  |  |

### IQGAP1 – phosphoinositide association is regulated by EGF stimulation

Fluorescently labeled PH-domains have been shown to accurately report the cellular distribution of PI(4,5)P_2_ and/or PI(3,4,5)P_3_ pools [22, 23]. The PH domain of PLCδ1 binds specifically to PI(4,5)P_2_ [24–26], while the PH domains of Akt1, Btk1 [25] and Akt2 [27] are sensors for PI(3,4,5)P_3_. For the eGFP-mCherry FRET pair, the distance at which half of the excitation energy of eGFP is transferred to mCherry is about 5.4 nm [28]. Here, we observe the interactions of IQGAP1 with fluorescently tagged pleckstrin homology (PH) domains of several proteins that bind to different phosphoinositides using FLIM-FRET. While the phosphoinositide lipid that binds the PH-domain is blocked from interacting with other PIP binding partners, this method is still successful in identifying pools of PIPs and hence, is useful to investigate protein/PIP interactions in a cellular context.

Using these sensors, We find an ~8% FRET-induced decrease in eGFP lifetime in HeLa cells that express both eGFP-IQGAP1 and mCherry PH-PLCδ1, confirming earlier results [16] that indicates an interaction between IQGAP1 and PI(4,5)P_2_. When the FRET and non-FRET populations are visualized separately, this interaction seems to be localized to plasma membrane and the nuclear membrane regions (**Fig. 3A**). This interaction is not affected by EGF or the PI3K antagonist, LY294002 (**Fig. 3B**). However, EGF stimulation enhances the interaction between eGFP-IQGAP1 and mCherry PH-Akt1 (which binds PI(3,4,5)P_3_), while treatment with LY294002 instead of EGF completely abrogates this interaction **(Fig. 3B)**. A significant FRET-induced lifetime decrease is also seen when co-expressing PH-Akt2 instead of PH-Akt1 but only after EGF treatment **(Fig. S4)**. These results strongly suggest that the interaction of IQGAP1 with PI(3,4,5)P_3_ is enhanced by EGF stimulation, in part due to increased levels of PI(3,4,5)P_3_.

**Figure 3:**
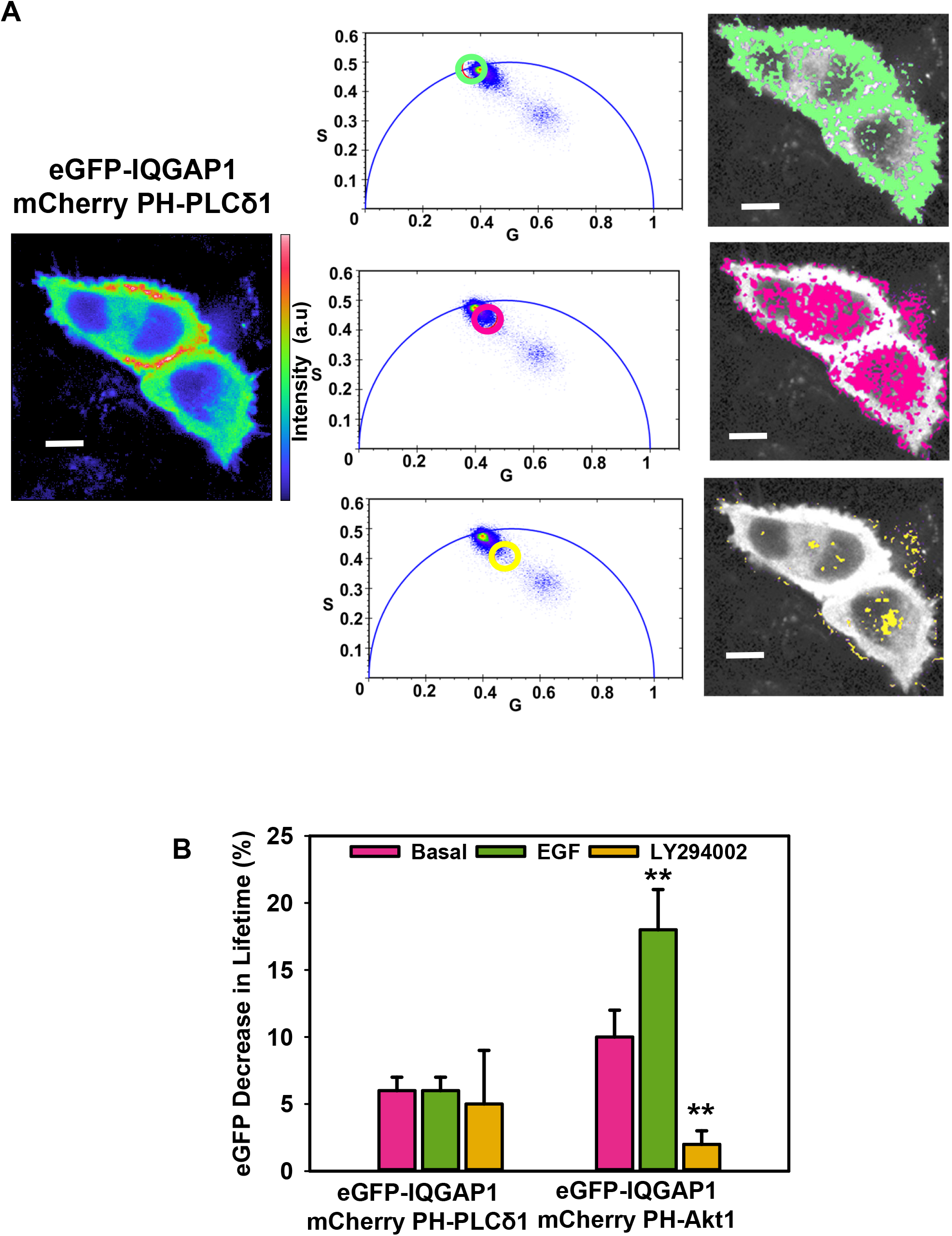
IQGAP1 associates with PI(4,5)P_2_ and PI(3,4,5)P_3_: A) The images of a representative HeLa cell co-expressing mCherry-PH-PLCδ1 and eGFP-IQGAP1 (left), where the pixels of distinct regions on phasor plots (center panels) are highlighted to show the distinct regions. The distinct regions on the phasor plots are represented by a green circle indicating higher (non-FRET) lifetimes (lifetime center = 2.55 ns) or by magenta or yellow circles indicating shortened lifetimes due to FRET (magenta lifetime center= 2.00 ns, yellow lifetime center= 1.60 ns). The pixels underlying these circles are false colored and overlaid on grayscale cell images (right panels). B) Expression of eGFP-IQGAP1 with both mCherry-PH PLCδ1 and mCherry-PH Akt1 in HeLa cells results in decreased eGFP IQGAP1 lifetimes due to FRET. For IQGAP1-PH-Akt1 interactions, the FLIM-FRET was significantly increased by PI3K agonist EGF (100 ng/ml) and decreased by PI3K antagonist LY294002 (1 μM) for (n≥4, **=p <0.001 from basal). In contrast, IQGAP1-PH-PLCδ1 interactions are not affected by EGF or LY294002. Error bars denote standard deviation. Scale bar= 10μm

### EGFR – phosphoinositide association is mediated by IQGAP1

IQGAP1 binds to the EGF receptor (EGFR), which is an upstream activator of the PI3K/Akt pathway [29, 30]. EGFR activity including its activation and cellular trafficking, are also known to be modulated by PI(4,5)P_2_ [31] and PI(3,4,5)P_3_ [32] levels. Here, we see FRET between eGFP-EGFR and mCherry PH-Akt1 (**Fig. 4A**) is significantly increased upon stimulation with EGF possibly due to a spike in PI(3,4,5)P_3_ levels. This change can also be visualized by the increase in high FRET populations along the membrane (**Fig. 4B**) However, downregulation of IQGAP1 eliminated this EGF-mediated increase in the EGFR-PH-Akt1 association **(Fig. 4C)**. These findings suggest that IQGAP1 links EGFR and PI(3,4,5)P_3_ by sequestering the receptor in the scaffolding complex. These results indicate that IQGAP1 scaffolds EGFR, its downstream effectors (PIPKI and PI3K kinases) and the substrates (phosphoinositides) on the membrane and thus IQGAP1-PIPKI interactions may act as the hub for this scaffold **(Fig. 4D)**.

**Figure 4:**
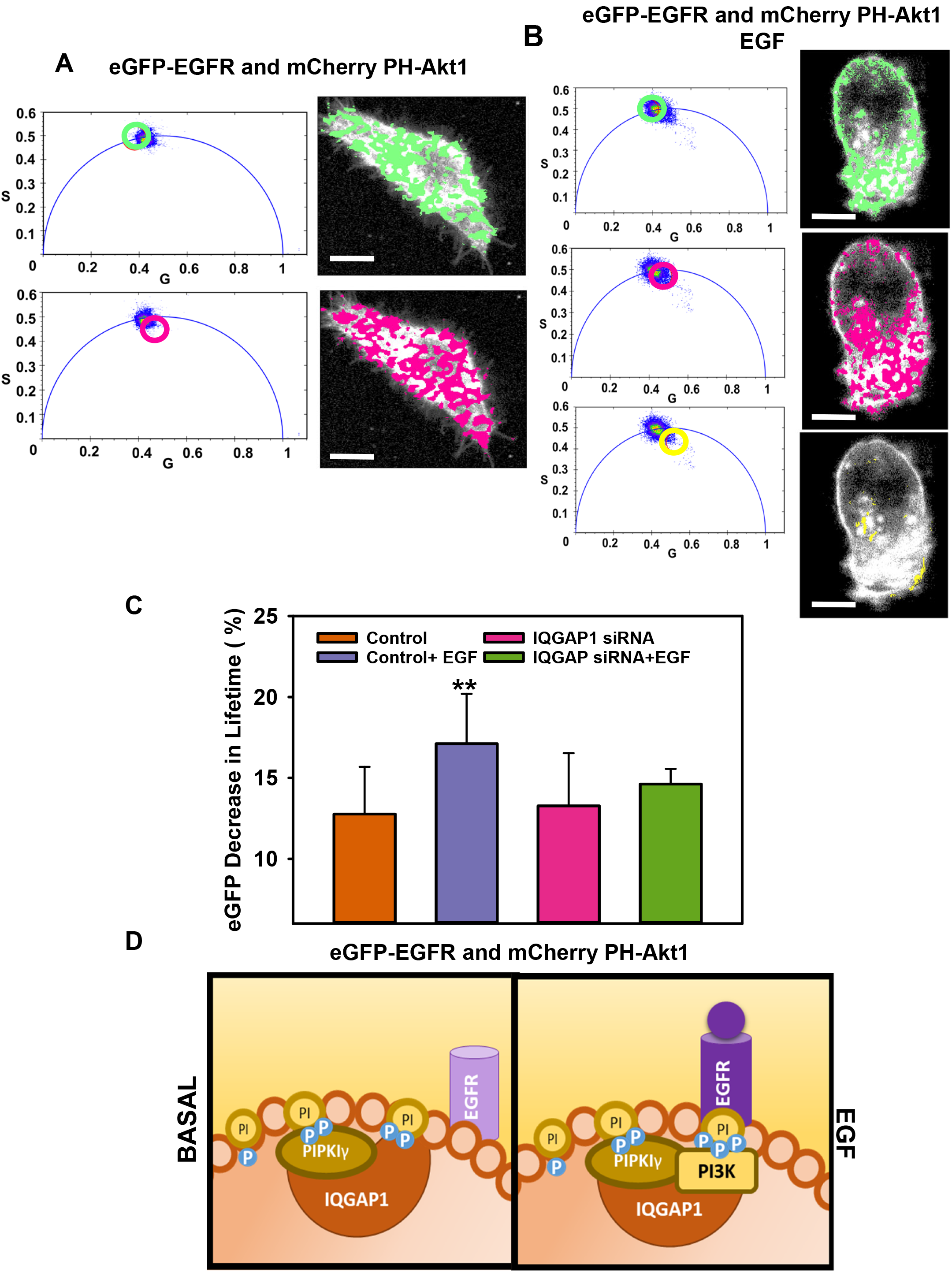
EGFR – phosphoinositide association is mediated by IQGAP1: A) The images of a representative HeLa cell co-expressing eGFP-EGFR and mCherry PH-Akt1,where the pixels of distinct regions on phasor plots (left panels) are highlighted to show the distinct regions. The distinct regions on the phasor plots are represented by green circle indicating higher (non-FRET) lifetimes (lifetime center = 2.55 ns) or by a magenta circle indicating shortened lifetimes (lifetime center= 2.00 ns). The pixels underlying these circles are false colored and overlaid on grayscale cell images (right panels). B) The images of a representative HeLa cell co-expressing eGFP-EGFR and mCherry PH-Akt1 stimulated by EGF (100 ng/ml) for 1 hour, where the pixels of distinct regions on phasor plots (left panels) are highlighted to show the distinct regions. The distinct regions on the phasor plots are represented by green circle indicating higher (non-FRET) lifetimes (lifetime center = 2.55 ns) or by magenta or yellow circles indicating shortened lifetimes (magenta lifetime center= 2.00 ns, yellow lifetime center= 1.60 ns). The pixels underlying these circles are false colored and overlaid on grayscale cell images (right panels). C) There is FLIM-FRET between eGFP-EGFR and mCherry-PH Akt1 that is significantly increased upon EGF stimulation (100 ng/ml). This EGF-mediated increase in FRET is abrogated upon downregulation of IQGAP1 using siRNA. All eGFP lifetimes are statistically lower compared to eGFP lifetimes in cells expressing only eGFP-EGFR (n≥4, **=p <0.001 from basal). These results indicate that IQGAP1 scaffolds a pathway at the plasma membrane, including its receptor (EGFR), downstream effectors (PIPKI and PI3K kinases) and the substrates (phosphoinositides). Since this scaffold is predominantly localized to the plasma membrane, IQGAP1-PIPKI interactions may act as the lynchpin for this scaffold as represented by a schematic (D). Error bars denote standard deviation.

### IQGAP1 forms complexes in response to pro-migratory signals

Consistent with our results of IQGAP’s increased interactions with PI(3,4,5)P_3_ upon stimulation with EGF, we observe a gradual but significant increase in eGFP-IQGAP1 intensity that is localized to the plasma membrane including cell-cell junctions **(Fig. 5A, B)**. The potential of IQGAP1 to oligomerize has been discussed [33, 34]. To investigate if the increase in eGFP-IQGAP1 fluorescence intensity also involves IQGAP1 oligomer formation, we used a number & brightness (N&B) method (described in *Experimental Procedures*). Here, we measured brightness (B) values to monitor eGFP-IQGAP1 complexes, where higher B values indicate the presence of oligomers **(Fig. 5C, D)**. After background correction and calibration, the B value corresponding to eGFP-IQGAP1 in HeLa cells increased slightly from 1.02 to 1.07 one hour after stimulation with EGF. However, the proportion of oligomeric species denoted by pixels with a B value above 1.5 that corresponds to the B values for dimers and greater almost doubled from 6% to about 12% upon treatment with EGF [35]. Although this is a small fraction of the total number of pixels, its significance is demonstrated by the predominance of the localization of this high-brightness population on the plasma membrane and cell-cell junctions **(Fig. 5C, D, E)**. These results indicate formation on larger IQGAP1 clusters including IQGAP1 oligomers at the plasma membrane population upon EGF stimulation **(Fig. 5F)**

**Figure 5:**
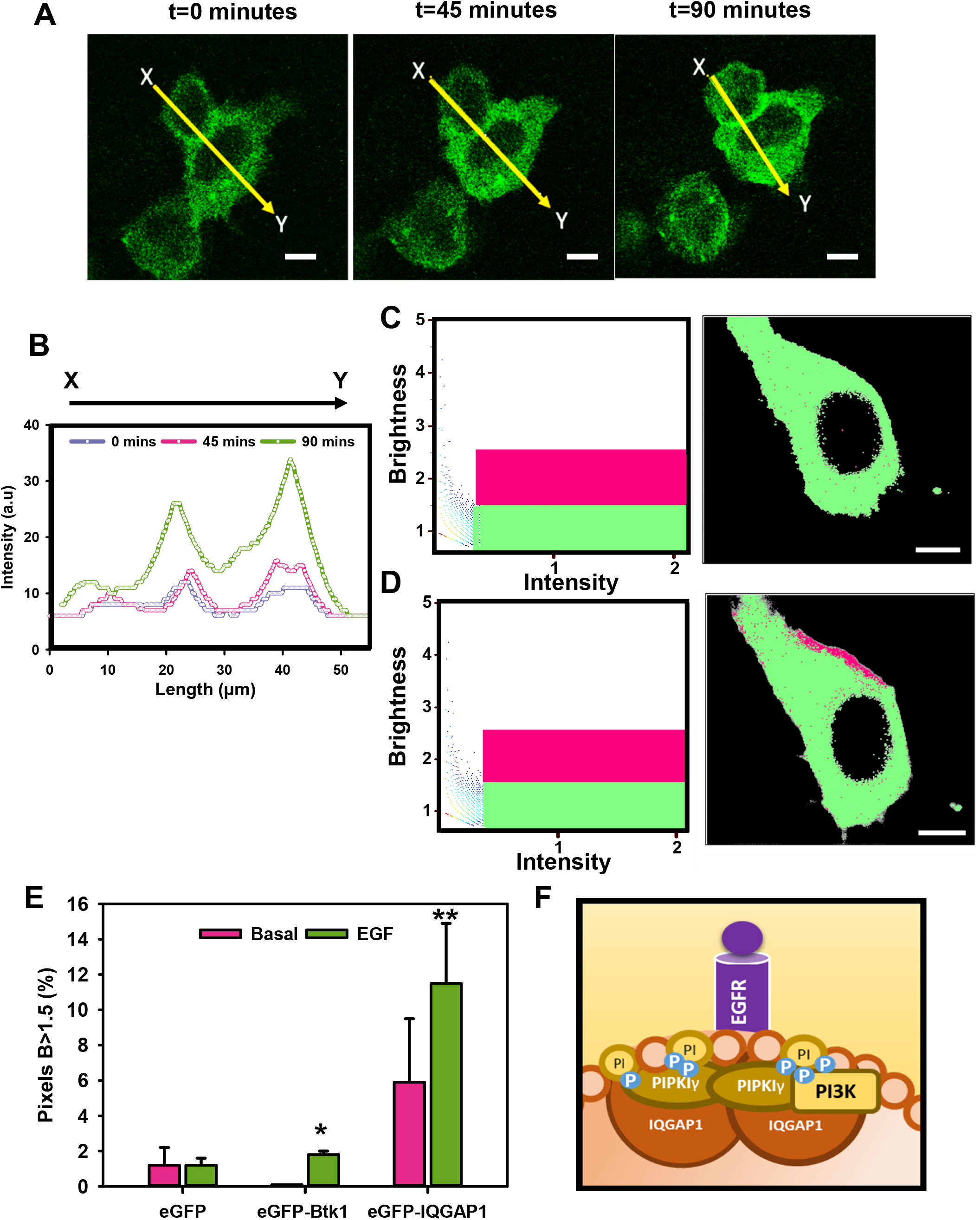
IQGAP1 forms membrane complexes in response to EGF: A)eGFP intensity is quantified along a 50 μm line from regions X to Y immediately, 45 minutes and 90 minutes after stimulation with 100 ng/ml EGF in HeLa cells expressing eGFP-IQGAP1.B) Upon quantification, these images in 3A show that eGFP-IQGAP1 levels increase along the plasma membrane in response to EGF. C)N&B results are plotted using a Brightness vs Intensity plot, where individual pixels of the HeLa cell images expressing eGFP-IQGAP1 are highlighted either in a green box (B<1.5) or a magenta box (B>1.5) (left panels). The distribution of these highlighted pixels can be seen overlaid (right panels) on a representative unstimulated cell. D) A Brightness vs Intensity plot, where individual pixels of HeLa cell images that is stimulated by EGF (100 ng/ml for 1 hour) expressing eGFP-IQGAP1 are highlighted either in a green box (B<1.5) or a magenta box (B>1.5) (left panels). The distribution of these highlighted pixels can be seen overlaid (right panels) on a representative cell This EGF-stimulated cell shows localization of IQGAP1 clusters at the plasma membrane. E) There is a statistically significant increase in the number of pixels having higher B values (magenta pixels, B >1. 5) that represent the oligomeric species or clusters of eGFP-IQGAP1 after the cells are stimulated with EGF compared to basal levels. (n≥5, *= p=0.03, **=p<0.001 from basal). For more details, see *Experimental Methods*. These results indicate that IQGAP1 clusters form along the membrane upon EGF stimulation, as represented by a schematic (F). Error bars denote standard deviation. *Scale bar= 10μm*

### IQGAP1 regulates β-actin complex formation in response to pro-migratory signals

Stimulation by pro-migratory agonists such as EGF results in periods of intense cytoskeletal remodeling [36]. These periods are characterized by increased actin polymerization facilitated by the Arp2/3 and VASP complex providing the force for lamellipodial protrusion that is characterized by the formation of actin clusters [37]. IQGAP1 is known to bind to and mediate the activities of actin as well as several other actin-binding proteins including Arp2/3 and VASP complex **(Table 1, Fig. S5)**. [38, 39]. PIPKIγ is also known to bind to actin and has been observed to be critical for actin cytoskeletal reorganization [40]. Both eGFP-IQGAP1 **(Fig. 6A)** and eGFP-PIPKIγ *(not shown)* were shown to strongly bind to mCherry β-actin in HeLa cells when quantified by FLIM–FRET corroborating these previous studies. Using N&B analysis, we detected a few mCherry-β-actin clusters primarily in the interior of the cells **(Fig. 6B)**. EGF stimulation resulted in an increase in the amounts and size of membrane clusters **(Fig. 6C)**. Knockdown of IQGAP1 resulted in a significant decrease (p<0.05) of these clusters. However, the amounts of β-actin clusters were partially restored by EGF treatment **(Fig. 6D)**, indicating that IQGAP1 is only partly responsible for β-actin complex formation. It is also possible that IQGAP-phosphoinositide interactions also have a similar effect on other cytoskeletal components that bind to IQGAP1, PI(4,5)P_2_ and PI3K**(Fig. S5)**.

**Figure 6:**
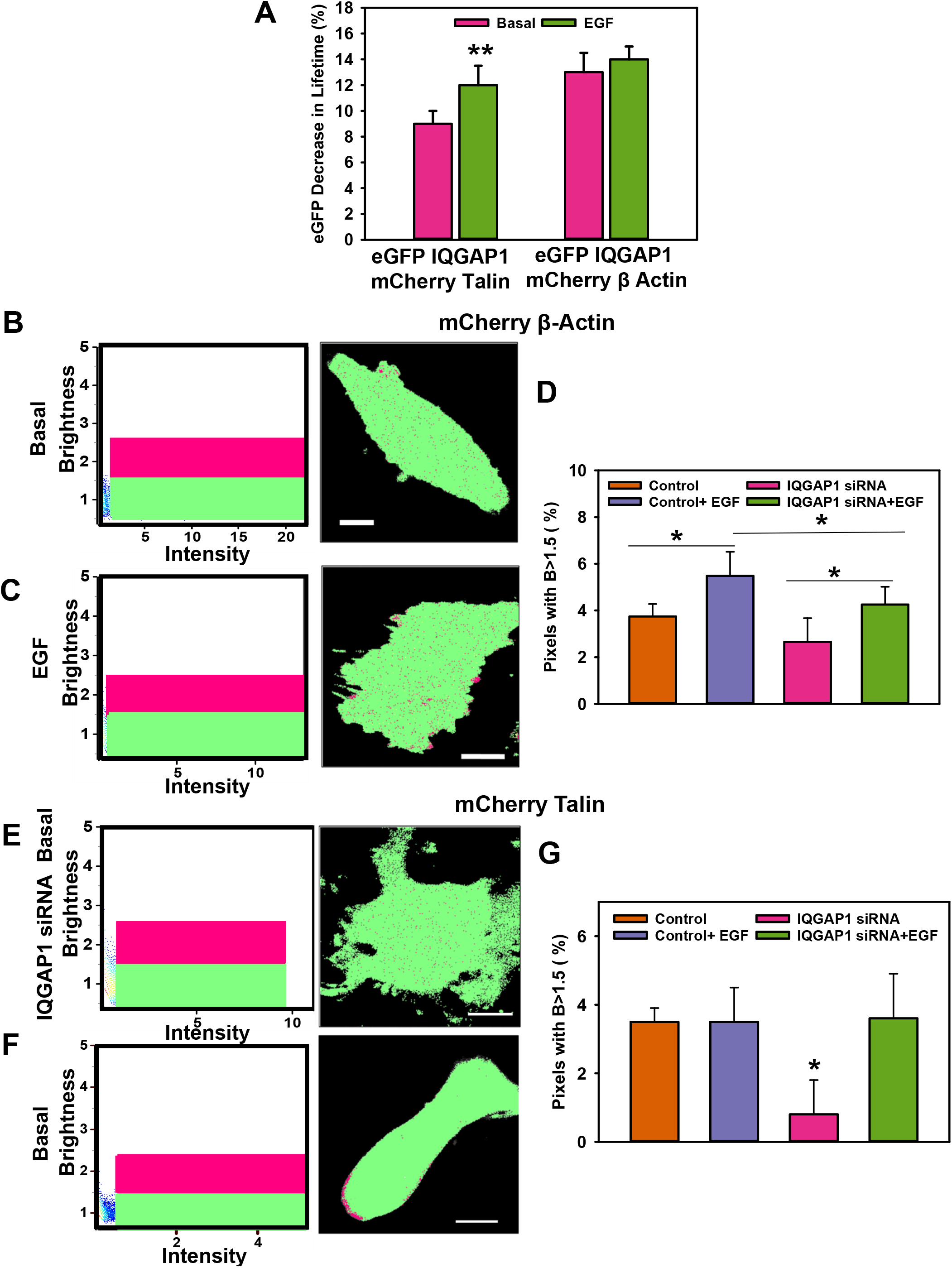
IQGAP1 interacts with β-actin and talin and mediates clustering: A) There is a decrease in fluorescence lifetime of eGFP-IQGAP1 due to FRET between eGFP-IQGAP1 and mCherry-talin and mCherry-β-actin in HeLa cells (n≥4, **= p<0.001). The interaction between IQGAP1 and talin is further enhanced by EGF stimulation (100 ng/ml). In all cases, eGFP lifetimes are significantly different from eGFP lifetimes in cells expressing eGFP-IQGAP1 alone (p<0.001). B,C) N&B results are plotted using a Brightness vs Intensity plot, where individual pixels of the images of HeLa cells expressing mCherry-β-actin are highlighted either in a green box (B<1.5) or a magenta box (B>1.5) (left panels). The distribution of these highlighted pixels can be seen overlaid (right panels) on a representative unstimulated cell. C) A Brightness vs Intensity plot, where individual pixels of the images of HeLa cells expressing mCherry-β-actin are highlighted either in a green box (B<1.5) or a magenta box (B>1.5) (left panels). The distribution of these highlighted pixels can be seen overlaid (right panels) on a representative cell that is stimulated by EGF (100 ng/ml) for 1 hour. D) A statistically larger number of pixels with higher B values (magenta pixels, B>1.5) that represent oligomeric species or clusters of mCherry-β-actin are seen after the cells are stimulated with EGF (*= p<0.001, n≥4 from basal). A similar but muted response is seen when IQGAP1 expression is downregulated using siRNA (*= p<0.05 from basal, n≥4) but this increase is statistically lower than the amount of actin clusters in stimulated control cells (p<0.05). This indicates that IQGAP1 is partly responsible for the formation of actin clusters in cells. E) A Brightness vs Intensity plot, where individual pixels of the HeLa cell images expressing mCherry-talin are highlighted either in a green box (B<1.5) or a magenta box (B>1.5) (left panels). The distribution of these highlighted pixels can be seen overlaid (right panels) on a representative unstimulated cell. F) A Brightness vs Intensity plot, where individual pixels of the HeLa cell images expressing mCherry-talin are highlighted either in a green box (B<1.5) or a magenta box (B>1.5) (left panels). The distribution of these highlighted pixels can be seen overlaid (right panels) on a representative cell that is stimulated by EGF (100 ng/ml) for 1 hour. G) mCherry-talin oligomeric species are significantly reduced in HeLa cells when treated with IQGAP1 siRNA (p<0.05 from control, n≥4), but the number of these talin clusters can be restored upon treatment with EGF (p<0.05 from basal, no difference from control). This indicates that IQGAP1 partly mediates the formation of talin clusters in cells. Scrambled (non-specific) siRNA treatment is used for the control groups. Error bars denote standard deviation. *Scale bar= 10μm*

IQGAP1 has been found to form complexes with Cdc42, Guanine Nucleotide exchange factor FGD6, the Rho GTPase-activating protein ARHGAP10, filamin and talin close to focal adhesions[41]. Talin-PI(4,5)P_2_ mediate the activation and the function of integrins [42], including the formation of talin-integrin clusters [43]. eGFP-IQGAP1 strongly binds to mCherry-talin in HeLa cells when quantified by FLIM–FRET and is slightly enhanced by EGF stimulation **(Fig. 6A)** However. EGF has no effect on the formation of talin clusters on basal cells **(Fig. 6G)**. Knockdown of IQGAP1 decreases the number of talin clusters **(Fig. 6E)**, but EGF treatment restores the levels of talin clusters in these cells **(Fig. 6F)**. Similarly, IQGAP-phosphoinositide interactions likely have a similar effect on the several focal adhesion complex proteins that are known to bind to PI(4,5)P_2_, IQGAP1 and PI3K **(Fig. S5)**. These indicate the potential of the IQGAP1-phosphoinositide interactions in scaffolding the focal adhesion proteins to other components of the cell migration signaling pathway.

### IQGAP1 controls PIPKIγ interactions with its binding partners Talin and Cdc42

PI(4,5)P_2_ and PIPKIγ have been found to play a major role in cell migration, especially by modulating actin reorganization and focal adhesion[40, 44], in conjugation with IQGAP1 [16, 17, 41]. A proteomics screen of proteins that were pulled down with endogenous IQGAP1 in HeLa cells revealed that IQGAP1 binds to several proteins that are known to be involved in these roles **(Table 1, Fig. S5)**. Interestingly, almost all of these proteins have already been found to interact with PI(4,5)P_2_, PIPKIγ or PI(3,4,5)P_3_ [45–47]. PIPKIγ is the only PIPKI isoform known to bind to talin [46]. Using FLIM-FRET, we see that eGFP-PIPKIγ interacts with mCherry-talin only when the cells are stimulated by EGF. This EGF-mediated increase in FRET (decrease in donor lifetime) was completely abrogated when IQGAP1 is knocked down, indicating that the PIPKIγ-talin interaction is probably scaffolded by IQGAP1 **(Fig. 7A)**.

**Figure 7:**
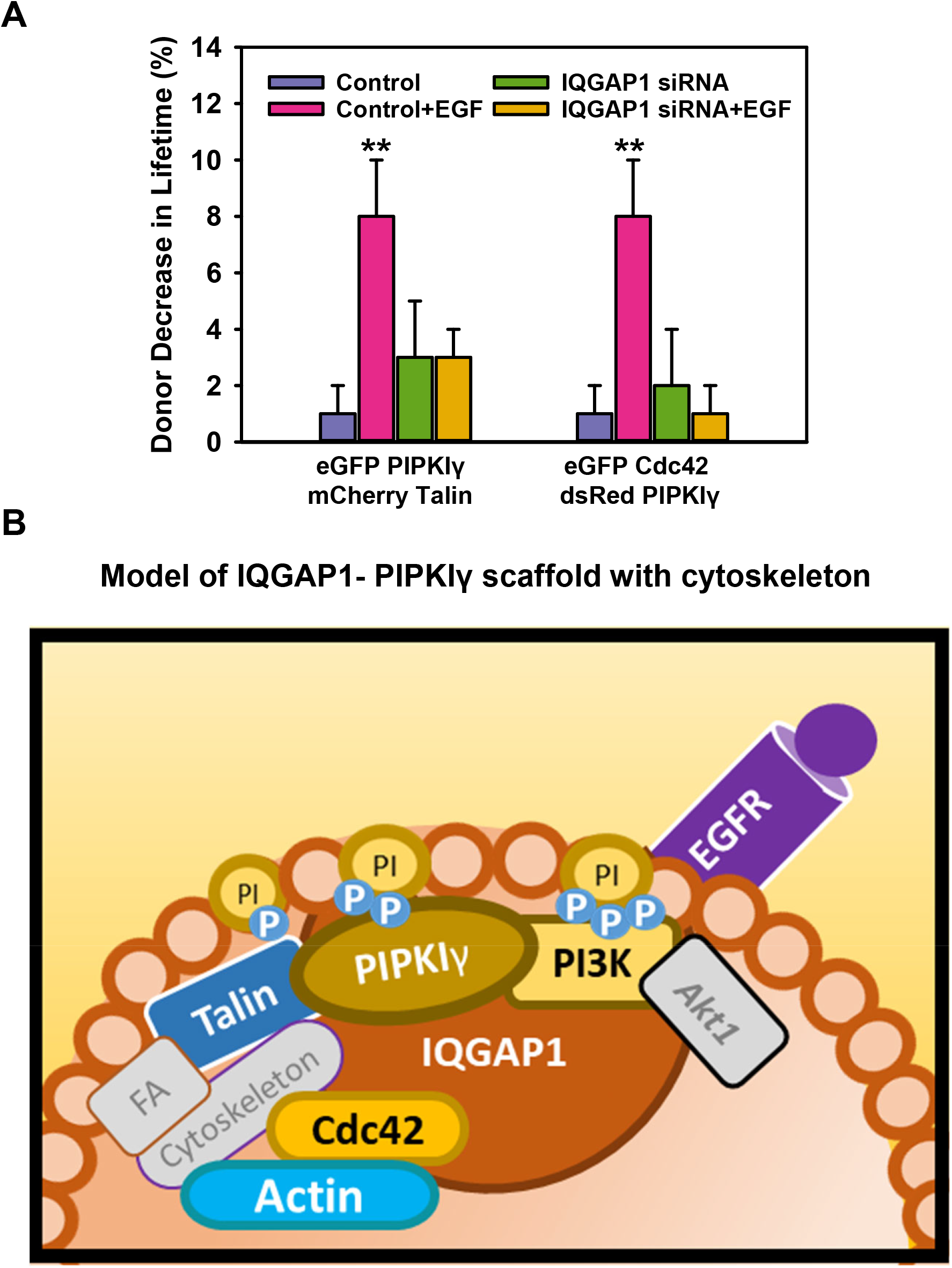
Interactions of PIPKIγ with talin and Cdc42 require IQGAP1: A) eGFP lifetime decreases due to FRET in HeLa cells expressing both eGFP-PIPKIγ and mCherry-talin only when stimulated with EGF (100 ng/ml) (n≥4, **=p<0.001 from basal). These EGF-mediated interactions are abrogated by IQGAP1 downregulation (n ≥ 4), indicating that IQGAP1 is required for PIPKIγ and talin interactions. A similar effect is seen in HeLa cells expressing both eGFP-Cdc42 and dsRed-PIPKIγ where FRET is seen only upon EGF (100 ng/ml) stimulation but not when IQGAP1 is downregulated (n≥4, **= p<0.001 from basal), indicating that IQGAP1 is required for PIPKIγ and Cdc42 interactions. Scrambled (non-specific) siRNA treatment is used for the control groups. Error bars denote standard deviation. B) IQGAP1 forms clusters along the plasma membrane to form a nexus between phosphoinositides, phosphoinositide kinases, receptors, focal adhesion proteins and cytoskeleton that promotes directionality and motility. Because PIPKIγ is essential for IQGAP1 localization at the membrane, our model indicates that IQGAP1-PIPKIγ interactions form the cornerstone for IQGAP1 scaffolding. Extracellular and intracellular signals such as EGF modulate this complex, which in turn regulates the movement of a cell by integrating cell migration, cytoskeletal reorganization and focal adhesion formation.

Cdc42 is a major initiator of the formation of the leading-edge of a moving cell. Cdc42 and other small GTPases RhoA and Rac1 are known to stimulate PIPKI activity and elevate the cellular levels of PI(4,5)P_2_ [48]. However, Cdc42 is not known to physically associate with any PIPKI isoforms unlike Rac1 and RhoA [49]. Interestingly, our studies reveal that mCherry-PIPKIγ exhibits FLIM-FRET with eGFP-Cdc42 only when the cells are stimulated by EGF. However, this EGF-mediated interaction is eliminated when IQGAP1 is knocked down using siRNA, indicating that a PIPKIγ–Cdc42 interaction is scaffolded by IQGAP1 when cells are stimulated with EGF **(Fig. 7A)**. These results indicate that IQGAP1 provides a physical link between phosphoinositides (through PIPKIγ), focal adhesion formation (through talin) and cytoskeletal reorganization (through Cdc42) upon EGF stimulation

## Discussion

Here, we present evidence that IQGAP1 dynamically interacts with phosphoinositide kinases and phosphoinositides to mediate phosphoinositide kinase interactions with its other binding partners as summarized in **Fig. 7B**. We utilize a variety of techniques including immunofluorescence, mass spectrometry and biophysical techniques to study these interactions. Specifically, FLIM-FRET studies in contrast to previous studies that used western blotting, immunoprecipitation and cell fixation, allow us to visualize protein-protein interactions in living cells

We see that IQGAP1’s interactions with PIPKI are stable in the presence of various agonists or antagonists, and are found in multiple cell lines. We also find that IQGAP1 promotes close localization of PI3K subunits and PIPKIγ. Since these kinases are not known to bind to each other, we show that IQGAP1 acts as a scaffold that enables their close proximity, giving credence to the idea that IQGAP1 promotes PI(3,4,5)P_3_ generation by scaffolding the phosphoinositide kinases close to phosphoinositides [17]. Therefore, our results that show an increase in IQGAP1 interactions with the PI(3,4,5)P_3_ sensors PH-Akt1 and PH-Akt2 in the presence of EGF indicates that IQGAP1 localizes to the sites of PI(3,4,5)P_3_ synthesis.

The signaling cascade that begins with PI3K generation of PI(3,4,5)P_3_ results in the activation of Akt and PDK1, which in turn regulates several downstream partners [50]. This signaling pathway regulates cell functions that include cell migration [1, 2]. Several of these downstream components of the PI3K/Akt pathway are also known to bind to IQGAP1 [51] (**Table 1**) (**Fig. S2C**). Physiologically, EGFR is one of the upstream activators of the PI3K/Akt pathway [29]. IQGAP1 is also known to bind to EGFR, where it modulates EGFR activity induced by EGF [30]. Here, we show that EGFR localizes to the sites of PI(3,4,5)P_3_ synthesis and that this localization is mediated by IQGAP1. Hence we believe that our results demonstrate the existence of a complex assembled by IQGAP1-PIPKIγ interactions that integrates the PI3K/Akt signaling pathways at the site of PI(3,4,5)P_3_ generation.

Dysregulation of the PI3K/Akt signaling pathway have been linked to enhanced cell migration and motility in various diseases such as cancer [1, 2]. IQGAP1 is upregulated in several different types of cancers [52], including hepatocellular [53, 54], prostate [55], glioma [56] and breast [57] cancers. In most studies, perturbing IQGAP1 levels by upregulation or downregulation has a corresponding effect on cell migration in a variety of normal and cancer cell lines [55–58]. It was also seen that IQGAP1 promotes cell migration in an interdependent manner with PIPKIγ. In various cancer cells, overexpression of either protein led to increased cell migration that was eliminated by knocking down the other protein [16]. Targeting and inhibiting the binding of PIPKIs and PI3K to the IQ and WW domains of IQGAP1 using peptide inhibitors reduced the number of viable cells in breast cancer cell lines with little or no effect on survival of normal cells [17]. In the context of these studies, IQGAP1-PIPKIγ scaffolds around regions of PI(3,4,5)P_3_ synthesis on the membrane that we observe could be a factor in these dysregulated cell migration conditions by scaffolding the pro-migratory PI3K/Akt signaling pathway.

EGF stimulation of HeLa cells was shown to not affect IQGAP1’s association with actin but enhances IQGAP1’s interaction with talin. However, we observe a striking increase in GFP-IQGAP1 localization along the plasma membrane and at cell-cell junctions upon stimulation. These findings highlight IQGAP1 moving closer to the membrane when the EGFR receptor is stimulated, dragging along actin. We also note an increase in the number of IQGAP1 clusters in response to EGF stimulation, potentially indicating an increase in the number of scaffolds or increasing the size of the scaffolds. Previous studies have shown that IQGAP1 is capable of dimerization [33] in a process mediated by Cdc42 [34] that may play a role in actin polymerization [59] [60]. Actin is known to form clusters during periods of intense cytoskeletal remodeling with other actin binding proteins such as Arp2/3 [38, 39]. Actin polymerization increases with stimulation with pro-migratory agonists such as EGF [36]. We observe inhibition of new actin clusters in Hela cells, especially along the plasma membrane in response to EGF stimulation using N&B experiments. We also observed that inhibition of the formation of actin clusters upon the knockdown of IQGAP1. Our results indicate a strong role of IQGAP1 in formation of actin clusters that potentially stems from IQGAP1’s interaction with many proteins that affect actin cytoskeleton, including talin and Cdc42.

Cdc42 is a major initiator of the formation of the leading-edge of a moving cell which is a site of intense actin polymerization and cell polarity. Cdc42 and other small GTPases RhoA and Rac1 are known to stimulate PIPKI activity and elevate the cellular level of PI(4,5)P_2_ [48]. Cdc42 is also a well-established binding partner of IQGAP1 [34, 61, 62], and they are known to localize together as an actin polymerization cluster. However, unlike Rac1 and RhoA, Cdc42 is not known to physically associate with any PIPKI isoforms [49]. Therefore, the proximity between PIPKIγ and Cdc42 is probably mediated by IQGAP1 scaffolding. Cdc42 is required for PI(4,5)P_2_-induced actin polymerization [63], and both Cdc42 and PI(4,5)P_2_ together mediate actin nucleation by activating WASP which in turn activates actin regulatory proteins [47]. We note that several proteins involved in actin nucleation and actin polymerization also bind to IQGAP1 and PI3K (Fig. S5).

We also demonstrate that IQGAP1 plays a role in mediating PIPKIγ binding to the PI(4,5)P_2_-binding focal adhesion protein, talin. PIPKIγ targets IQGAP1 to the plasma membrane and therefore to phosphoinositides [16]. PIPKIγ is the only PIPKI isoform known to bind to talin [46]. As this PIPKIγ-talin binding is regulated by phosphorylation [64], it is possible that the kinases involved in this process are also scaffolded by IQGAP1. The interaction between PIPK1 and talin have shown to be responsible for spatially mediating PI(4,5)P_2_ levels near focal adhesions [46]. These localized PI(4,5)P_2_ pools have been hypothesized to play a major role in mediating actin polymerization by affecting actin-capping proteins and by mediating the directionality of actin polymerization along the extracellular migration by interacting with talin and vinculin [3, 11, 12]. This interaction between PI(4,5)P_2_ and talin also mediates the activation and the function of integrins [42], including the formation of talin-integrin clusters [43]. We observe talin and several focal adhesion proteins bind to IQGAP1 **(Fig. S5)**. We also see that the number of talin clusters in HeLa cells are significantly lowered upon knockdown of IQGAP1. Through talin and integrin, IQGAP1 and PIPKIγ form a signaling nexus that can transmit from the extracellular matrix to the cytoskeleton.

IQGAP1 forms protein complexes that regulate cytoskeletal dynamics in several different organisms [65]. Our results indicate that IQGAP1 plays a role in facilitating the interplay between PIPKI, PI(4,5)P_2_, actin, focal adhesion proteins and cdc42 mediating important cell processes such as cytoskeletal reorganization, actin polymerization, leading-edge formation and focal adhesions. These findings also imply that IQGAP1 may scaffold interactions between the other phosphoinositide-interacting binding partners listed in **Table 1** that include many other proteins involved in cytoskeletal reorganization, that in turn regulate cell migration [66]. While the IQGAP1 knockout mice are viable and fertile, they suffer from gastric hyperplasia, delayed differentiation of neural progenitors and increased pulmonary vascular leak and high cardiac pressure overload [67]. In addition, the interactions between small GTPases, phosphoinositides, and actin play a role in vesicular trafficking and pre-synaptic pathways. These phenotypes can potentially be attributed to anomalies in cell motility caused by the loss of IQGAP1 scaffolding. Additionally, knockout of both IQGAP1 and IQGAP_2_ reverses the high incidence of hepatocellular carcinoma observed in IQGAP2-/- mice [54].

In summary, IQGAP1-PIPKIγ interactions provide a scaffold where different signaling pathways that regulate the cytoskeleton converge and interact. By corralling the PI(4,5)P_2_ and PI(3,4,5)P_3_ machinery to the regulators of the cytoskeletal elements, IQGAP1 co-ordinates cytoskeletal rearrangements in response to various stimuli. This coordination allows protein complexes generated by IQGAP1 scaffolding to mediate cell motility, cytoskeletal reorganization, actin polymerization, leading-edge formation and focal adhesions. Thus, this crosstalk mediated by IQGAP1 and phosphoinositides provides for a well-regulated and polarized cell migration.

### Experimental Procedures

#### Materials

HeLa, HepG2A and NIH/3T3 cells were obtained from American Type Culture Collection (Manassas, VA, USA). Untagged eGFP and mCherry plasmids were obtained from Clontech (Mountain View, CA, USA). dsRed-PIPKIγ was a kind gift from Dr. Richard Anderson (University of Wisconsin, Madison, WI, USA). PI4K plasmids, eGFP-PH Btk1,eGFP-PH-PLCδ1, mCherry-PH-SidM were a kind gift from Dr. Tamas Balla (NIH, Bethesda, MD, USA), PI3K 110 plasmids were a kind gift from Dr. Jonathan Backer (Albert Einstein School of Medicine, Bronx, NY, USA) All other plasmids were obtained through AddGene (Boston, MA, USA) from the following investigators: eGFP-IQGAP1 (Dr. David Sacks), eGFP-PIPKIγ (Dr. Pietro de Camilli) eYFP-p85 PI3K (Dr. Lewis Cantley), mCherry PH-PLCδ1(Dr. Narasimhan Gautam), mCherry-PH-Akt1(Dr. Moritoshi Sato), mCherry-PH-Akt2 (Dr. Ivan Yudishkin), eGFP-Cdc42 (Dr. Gary Bokoch), mCherry Talin (Dr. Michael Davidson). On-Target Plus SmartPool IQGAP1-siRNA (GE Dharmacon, Lafayette, CO, USA) was used to knockdown IQGAP1 while the scrambled siRNA was obtained from Ambion (ThermoFisher Scientific, Waltham, MA, USA).

#### Cell Culture

HeLa, HepG2A and NIH 3T3 cells were incubated in high-glucose DMEM (ThermoFisher Scientific) supplemented with 5% fetal bovine serum (Atlanta Biologicals, Flowery Branch, GA, USA) at 37°C with 5% CO_2_. Transfection of plasmids and small interfering RNA (siRNA) was performed using Lipofectamine 3000 (ThermoFisher Scientific) in antibiotic-free medium as per the manufacturer’s instructions. For treatment with agonist, cells were cultured in a serum-free high-glucose DMEM for 2 hours, following which they were supplemented with EGF (100 ng/ml), isoproterenol (10 μM) or LY2942001 (1 μM) and incubated for at least 1 hour

#### Confocal Imaging/Immunofluorescence

Cells plated on MatTek chambers (MatTek, Ashland, MA, USA) were fixed using 3.7% formaldehyde and permeabilized with 0.2% nonyl phenoxypolyethoxylethanol (NP40) in phosphate buffered saline (PBS) for 5 min and then blocked in PBS containing 1% BSA for 1 h. If appropriate, cells were then incubated with the primary antibody diluted to 1:1000 for 1.5 hours at 37°C, followed by incubation with Alexa-labeled secondary antibody for 0.5 hours at the same temperature. Cells were washed with Tris buffered saline (TBS) buffer after the incubations. Images of the cells were obtained using a Zeiss LSM 510 confocal microscope, and were analyzed using Zeiss software and ImageJ [68] (National Institutes of Health, Bethesda, MD,USA). Live cell images were taken on the same microscope while the cells were incubated in a custom-built chamber at 37°C and 5% CO_2_.

#### Fluorescence lifetime imaging microscopy (FLIM)

FLIM is a method to measure Förster resonance energy transfer (FRET) between labeled proteins in cells. FRET reduces the fluorescence lifetime of a donor molecule due to transfer of energy to an acceptor. Because of its steep distance dependence in the low nanometer range, FRET transfer requires a direct physical interaction between the proteins to which the probes are attached. If the donor is excited with intensity modulated light, then the phase of the acceptor fluorescence will be shifted. The fluorescence lifetime can be calculated from these phase shifts. We can determine the phase and modulation for each individual pixel of the image and plot the phase shifts and modulation decreases on a phasor plot. The phasor plot is a combined graphical representation of all the raw FLIM data in a vector space. The phasor space is constructed by using two phasor vectors (G,S), where each component is represented as shown in Eq.1:

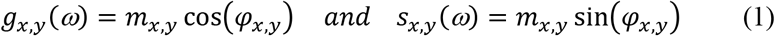

m_x,y_ and ϕ_x,y_ are the modulation ratio and the phase delay measured for a particular modulation frequency (ω) at a pixel location (x,y). For a single lifetime population, the values from all of the pixels will fall on the phasor arc with longer lifetimes displayed to the left and shorter lifetimes, to the right. Since FRET shortens donor lifetimes, FRET will manifest itself by moving the (g(ω),s(ω)) data point to the right. For a mixed population of donor molecules, some of which undergoing FRET, the resulting phasor points localize inside the phasor arc [69]. Distributions inside the phasor arc therefore indicate FRET transfer and, hence, association of the two labeled proteins. The lifetime of an individual pixel can be calculated by:

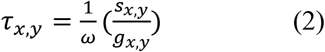

For our experiments, we used green fluorescent donors (such as eGFP, enhanced Green Fluorescent Protein) or yellow fluorescent donor eYFP with red fluorescent proteins such as mCherry and dsRed acting as the FRET acceptors.

FLIM was performed by acquiring images of live cells plated on MatTek chambers using a 2-photon MaiTai laser (Spectra-Physics, Santa Clara, CA, USA) (excitation 850 nm at 80 MHz) and a Nikon inverted confocal microscope in an ISS Alba System (Champaign, IL, USA).The Images were analyzed using ISS VistaVision and ImageJ software packages. Atto 425 fluorescent dye (t=3.61 ns) was used to calibrate the sample lifetimes.

#### Number and Brightness (N&B)

The Number and Brightness (N&B) analysis is a powerful tool that has been used previously to quantify graphically the aggregation state of diffusing proteins in living cells[35, 70-72]. N&B analysis can determine of the number (N) of diffusing particles within a given focal area and the intrinsic brightness (B) of each particle represented by pixels in an image and provides providing a map of brightness for every pixel using the following equation (3):

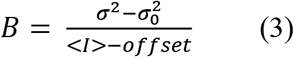

where I denotes the intensity of the signal, σ^2^ is the variance of the signal, σ_0_^2^ is the readout noise variance of the detection electronics and offset is the detector offset. The analysis has been described in detail earlier [35]. Higher variance in fluorescence is associated with higher-order oligomers. Therefore, the brightness vs intensity map can be used to determine the size of the aggregate at a given location as the brightness B can directly relate to the dimensions of the fluorescent molecules [70]. N&B studies were performed, by acquiring images of live cells plated on MatTek chambers (MatTek, MA, USA) using a 2-photon MaiTai laser (Spectra-Physics, Santa Clara, CA, USA), a Nikon inverted confocal microscope in an ISS Alba System (Champaign, IL,USA).The Images were analyzed using ISS VistaVision and SimFCS 4 (Irvine, CA, USA) software packages. The analysis has been described in more detail [71, 72]

#### Co-immunoprecipitation

HeLa cells were lysed with 500 ul of buffer containing 150 mM NaCl, 20 mM HEPES, 2 mM MgCl2, 5 mM 2-mercaptoethanol, 1 mM phenylmethylsulfonyl fluoride and cOmplete™ protease inhibitor cocktail tablet (Roche/Millipore Sigma, St. Louis, MA, USA). After pre-clearing non-specifically binding proteins by incubating lysate with 20 μl of Protein A beads (ThermoFisher Scientific), the lysate incubated with 5 μl of anti-IQGAP1 antibody (AbCam, Boston, MA, USA) overnight at 4°C after the removal of beads. Subsequently an additional 20 μl of Protein A beads was added to the mixture, which was then gently rotated for 4 hours at 4°C. The unbound proteins were separated from the beads, which were then washed twice with the lysis buffer. The bound proteins were then eluted from the beads in sample buffer at 95°C for 3 minutes and were analyzed using SDS-PAGE and Western blotting.

#### Mass Spectroscopy

The protein sample obtained from co-immunoprecipitation was run on an SDS-PAGE gel and stained using Coomassie blue. The pieces of gel containing the protein were cut into 1×1 mm pieces and placed in 1.5 mL tubes with 1mL of water for 30 min. This sample was sent to a University of Massachusetts Medical School Mass Spectroscopy Facility (Dr. John Leszyk), for performing the mass spectroscopy as described previously [73]. Raw data files were peak processed with Proteome Discoverer (version 2.1, ThermoFisher Scientific) prior to database searching with Mascot Server (version 2.5) against the Uniprot database. Search results were then loaded into the Scaffold Viewer (Proteome Software, Inc.) for peptide/ protein validation and label free quantitation. The data was further analyzed using the Database for Annotation, Visualization and Integrated Discovery (DAVID) [74] (NIH).

### Statistics

All data were analyzed using Sigmaplot (SysStat, San Jose, CA, USA) using Student’s t-test substituted by Mann-Whitney’s rank sum test if the data was not normally distributed. A difference between two groups was considered statistically significant only with a p-value below 0.05.

## Acknowledgments

We would like to acknowledge Drs. Richard Anderson (University of Wisconsin, Madison), Jonathan Backer (Albert Einstein School of Medicine) and Tamas Balla (NIH) for helpful discussion. We want to thank Dr. John Leszyk for his assistance with mass spectrometry, Dr. Qi Wen (WPI) for providing the NIH/3T3 cells, and Yemi Osayame, Christopher Dallarosa and Dr. Osama Garwain for their assistance. This work has been funded by NSF CHE 1508499 (AG) and NIH R01GM 68737 (SS).

