## Supplementary information for "IQGAP1 connects phosphoinositide signaling to cytoskeletal reorganization"

**Figure S1: Changes in lifetime do not occur for eGFP and mCherry lacking functional domains:** A) A false color image of a representative cell (left) shows untagged eGFP expressed in HeLa cells. A phasor plot (middle) shows that eGFP has a single fluorescent lifetime indicated by a homogenous population on the phasor arc. The pixels included in the green circle of the phasor plot (lifetime center= 2.56 ns) are false colored green and overlaid on a grayscale image of the cell (right), demonstrating a uniform lifetime throughout the cell. B) Average eGFP lifetime in HeLa cells transfected with untagged eGFP, eGFP-IQGAP1, eGFP-IQGAP1 with untagged mCherry, eGFP-PIPKIγ, or eGFP-PIPKIγ with untagged mCherry (n≥5, no significant differences between with and without untagged mCherry groups). Error bars denote standard deviation. *Scale bar= 10μm*

**Figure S2: IQGAP1 strongly associates with PIPKIγ**: A) Distribution of each individual pixel lifetime shifts due to FRET between eGFP-IQGAP1 and dsRed-PIPKIγ ($p<0.001$, Medians 2.35 and 2.15). This histogram analysis reaches the same conclusion as the phasor plot in Fig. 1. B) Coexpression of dsRed-PIPKIγ with eGFP-IQGAP1 significantly decreases eGFP lifetime ($p<0.001$)). Treatment with EGF (100 ng/ml) or with LY294002 (1 μM) does not significantly affect this interaction. C) eGFP-IQGAP1 shows evidence of FRET when co-expressed with dsRed-PIPKIγ in NIH3T3 and HepG2A cell lines, in addition to HeLa results in decreased eGFP IQGAP1 lifetimes (n≥5, $p<0.001$ from eGFP-IQGAP only). Error bars denote standard deviation.

**Figure S3: IQGAP1 interacts with PIPKIγ, PI3K, Akt1 but not PTEN, and mediates an association between PIPKIγ and PI3K p110α:** A) A representative image showing colocalization between eGFP- PIPKIγ and anti-IQGAP1 antibody tagged with Alexa 647 secondary antibody. B) A representative image showing colocalization between eGFP-IQGAP1 and anti-PI3K p110α antibody tagged with Alexa 647 secondary antibody).C) Co-localization quantified using Pearson's coefficient based on immunofluorescence studies show that endogenous IQGAP1 co-localizes with both eGFP and dsRed tagged PIPKIγ, Akt1 but not PTEN, while eGFP-IQGAP1 co-localizes with PI3K p110α. (Positive Control= 0.53 +/- 0.02, Negative Control= 0.004 +/-0.007). D) FLIM/FRET studies show that mCherry-Akt1 associates with eGFP-IQGAP1 in HeLa cells. The FRET in both cases is unchanged by EGF (100 ng/ml) stimulation. In both these cases, eGFP lifetimes are statistically lower than those in cells that expresses only eGFP-IQGAP ($p<0.001$, n≥5) E) FLIM/FRET studies show that dsRed-PIPKIγ associates with both YFP-PI3K p85 and emGFP- PI3K p110α at a basal stage in HeLa cells. EGF enhances FRET between PI3K p85 subunit and PIPKIγ but not between PI3K p110α and PIPKIγ, as demonstrated by a decrease in fluorescence lifetime in these cells (**= $p<0.001$). However, LY294002 treatment eliminates interaction between PIPKIγ and PI3K p110α but not with PI3K p85 (*= p= 0.02, n≥5). Error bars denote standard deviation.

**Figure S4: IQGAP1 associates with PI(4,5)$P_2$ and PI(3,4,5)$P_3$ but not PI(4P)**: A) eGFP-IQGAP1 demonstrates FRET when co-expressed with mCherry-PH-PLCδ1 (PI(4,5)$P_2$ sensor), mCherry-PH-Akt1 and mCherry-PH-Akt2 (PI(3,4,5)$P_3$ sensor) but not mCherry-PH-SidM (PI(4)P sensor). (n≥4, ** = p <0.001) in HeLa cells. Only the IQGAP1-PH-Akt1 and IQGAP1-PH-Akt2 interactions are altered by EGF, (p <0.001). eGFP lifetimes of all PH-PLCδ1 and PH-Akt1 significantly different ($p<0.001$) from eGFP-IQGAP1 the except IQGAP+PH PLCδ1+EGF. eGFP lifetimes in cells expressing both eGFP-IQGAP1 and mCherry PH Akt2 are significantly different from cells that express eGFP-IQGAP1 only upon EGF stimulation ($p<0.001$). Error bars denote standard deviation.

**Figure S4: IQGAP1 and PI3K bind to focal adhesion and cytoskeletal proteins**: A) A graph of several focal adhesion proteins that were co-immunoprecipated with anti-IQGAP1 and anti-PI3K p110a antibodies that denotes their percent coverage which is calculated by dividing the number of amino acids in all found peptides by the total number of amino acids in the entire protein sequence. The proteins that are known to bind to PI(4,5)$P_2$ are highlighted. B) A graph of several cytoskeletal proteins that were co-immunoprecipated with anti-IQGAP1 and anti-PI3K p110a antibodies that denotes their percent coverage which is calculated by dividing the number of amino acids in all found peptides by the total number of amino acids in the entire protein sequence. The proteins that are known to bind to PI(4,5)$P_2$ are highlighted.

### Supplementary Figure 1

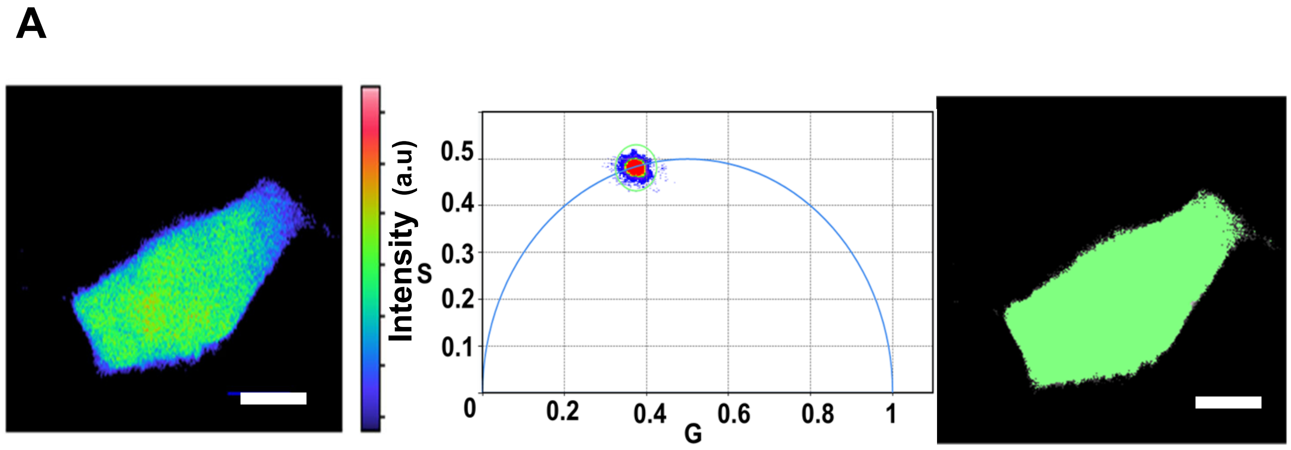


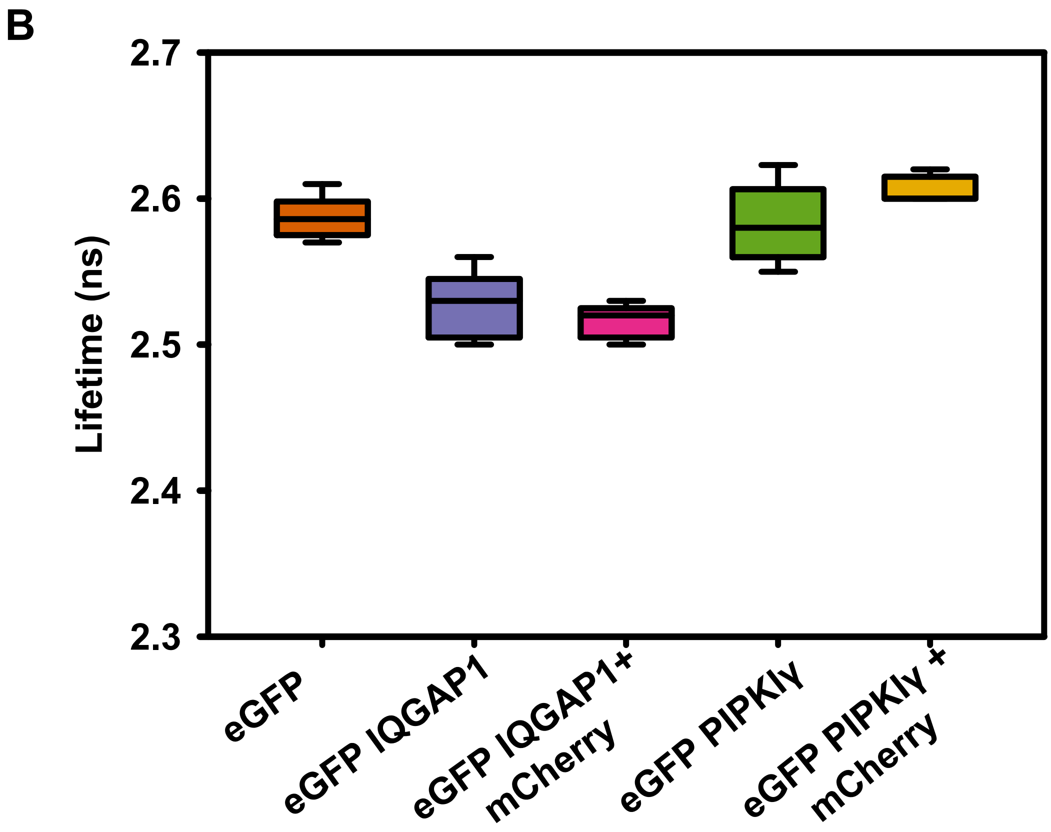

### Supplementary Figure 2

A

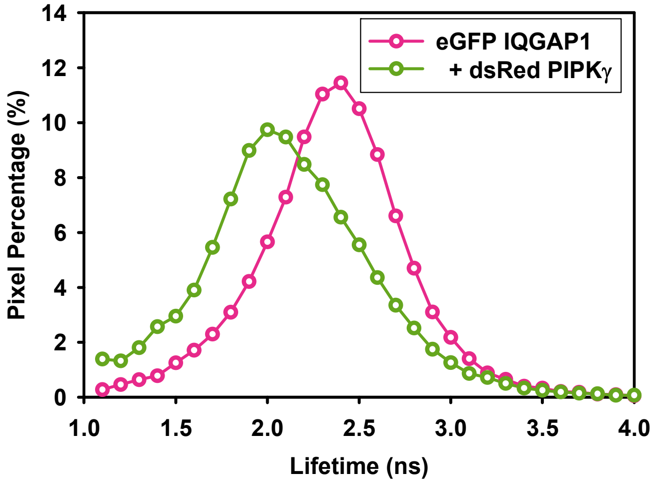


B

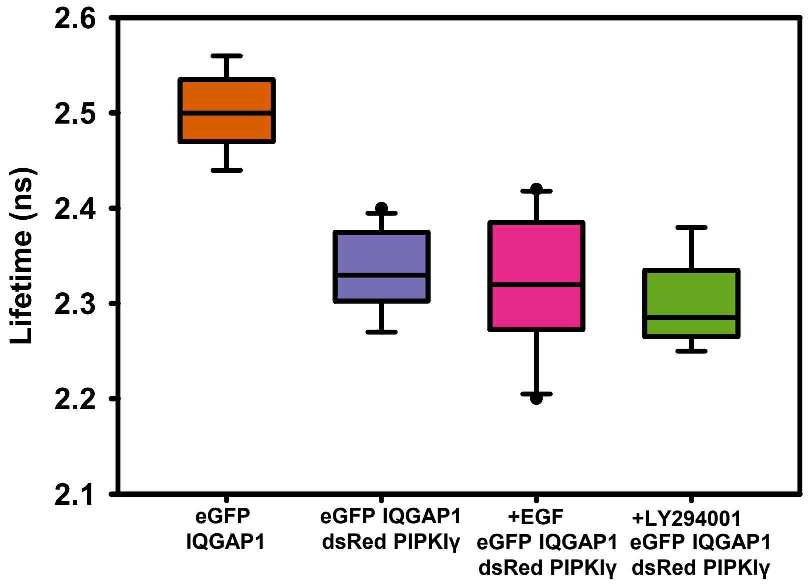


C

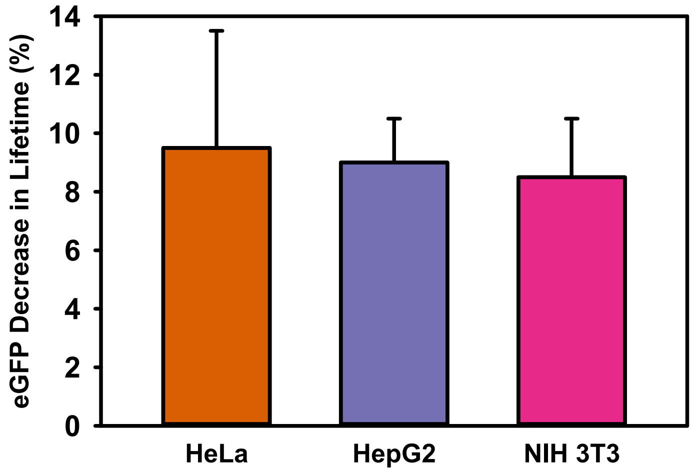

### Supplementary Figure 3

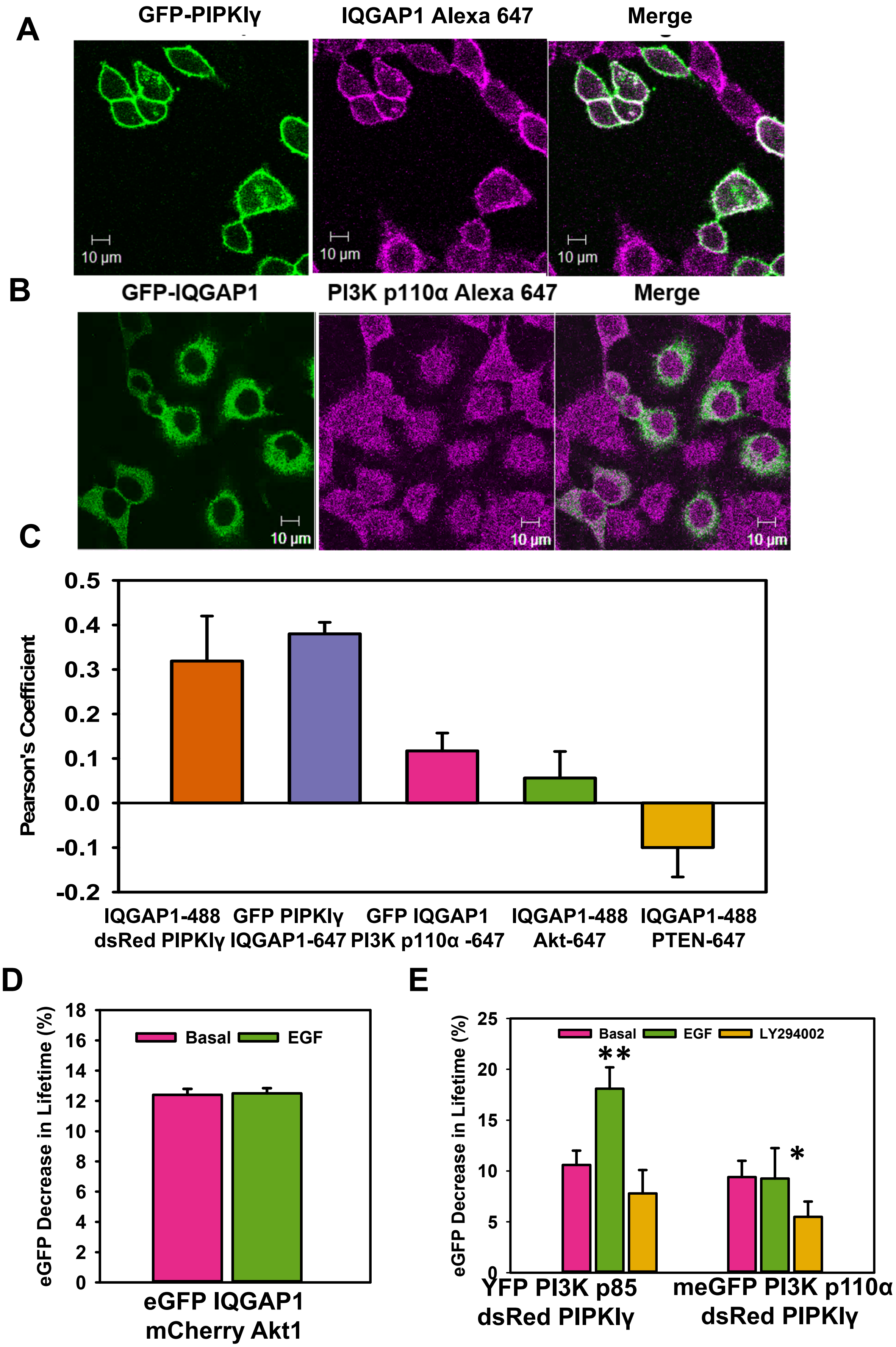

### Supplementary Figure 4

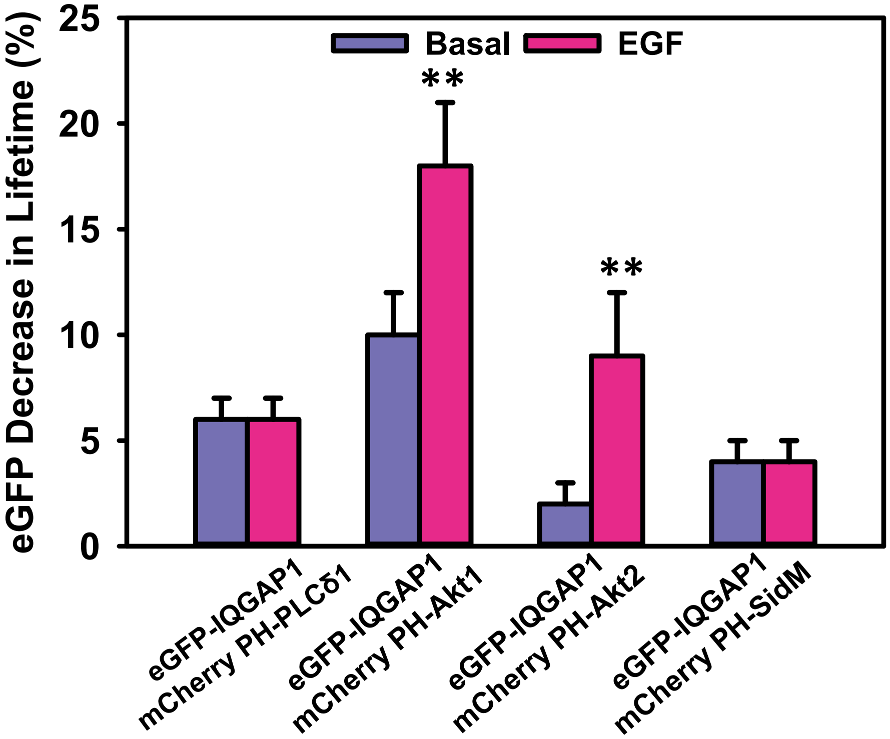

### Supplementary Figure 5

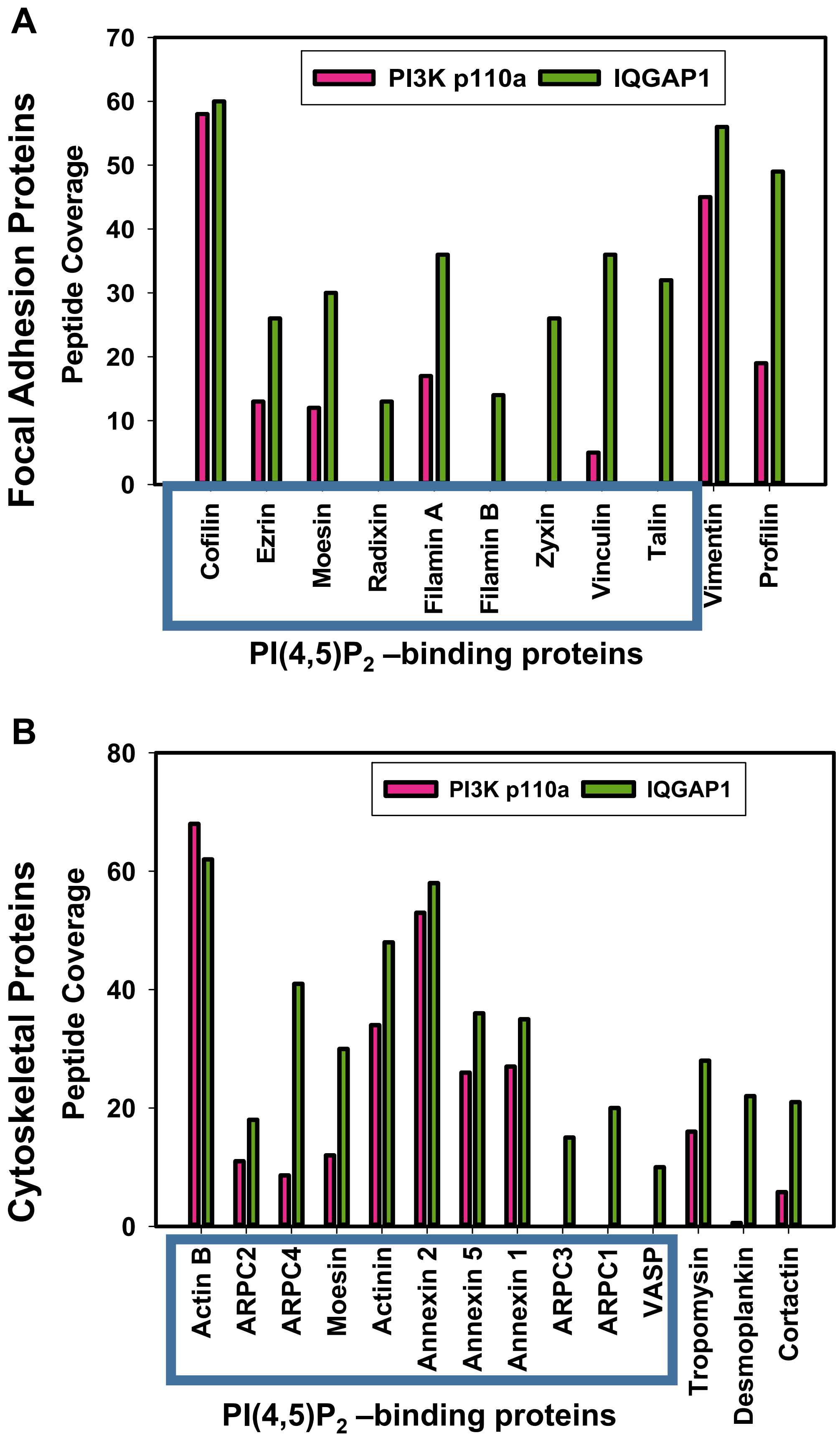
